# SEA: Simple Enrichment Analysis of motifs

**DOI:** 10.1101/2021.08.23.457422

**Authors:** Timothy L. Bailey, Charles E. Grant

## Abstract

Motif enrichment algorithms can identify known sequence motifs that are present to a statistically significant degree in DNA, RNA and protein sequences. Databases of such known motifs exist for DNA- and RNA-binding proteins, as well as for many functional protein motifs. The SEA (“Simple Enrichment Analysis”) algorithm presented here uses a simple, consistent approach for detecting the enrichment of motifs in DNA, RNA or protein sequences, as well as in sequences using user-defined alphabets. SEA can identify known motifs that are enriched in a single set of input sequences, and can also perform differential motif enrichment analysis when presented with an additional set of control sequences. Using *in vivo* DNA (ChIP-seq) data as input to SEA, and validating motifs with reference motifs derived from *in vitro* data, we show that SEA is is faster than three widely-used motif enrichment algorithms (AME, CentriMo and Pscan), while delivering comparable accuracy. We also show that, in contrast to other motif enrichment algorithms, SEA reports accurate estimates of statistical significance. SEA is easy to use via its web server at https://meme-suite.org, and is fully integrated with the widely-used MEME Suite of sequence analysis tools, which can be freely downloaded at the same web site for non-commercial use.

## 1 Introduction

Short, approximate sequence patterns (motifs) are known to encode functional signals in DNA, RNA and protein sequences. In genomic DNA, motifs capture the preferred binding sites of transcription factors (TFs), and numerous databases catalog the motifs associated with given TFs (e.g, the JASPAR database [3]). Similarly, databases of motifs bound by RNA-binding proteins (RBPs) have been compiled (e.g., the CISBP-RNA database [9]). The ELM database [6] contains short linear protein motifs annotated with the type of functional regions they encode in many proteins.

Motif enrichment analysis (MEA) algorithms can be used to determine if a group of sequences contains a statistically significant number of (approximate) matches to a given motif. When used in conjunction with databases of known motifs, MEA algorithms can suggest if the sequences may contain functional signals in common. For example, motif enrichment analysis of sets of gene promoters (e.g., of coexpressed genes) can identify which transcription factors may bind to a significant subset of the given promoters to regulate the expression of their associated genes. Another common application of MEA algorithms is to identify possible regulatory cofactors of a TF whose binding sites have been identified in a ChIP-seq experiment. TFs with motifs that differ significantly from the TF pulled down in the experiment, and that show significant enrichment in the ChIP-seq peak regions may regulate the expression of genes in concert with the ChIP-ed TF.

Several MEA algorithms are currently in wide use including Clover [4], Pscan [11], AME [8] and CentriMo [7]. All these algorithms use each candidate motif, encoded as a position weight matrix (PWM), to score the input sequences. The first three algorithms compare the scores of a set of (primary) sequences of interest with those of a control set of sequences. The fourth algorithm above (CentriMo) takes a different approach, comparing the scores of the central regions of the (primary) sequences with the scores of their flanks.

The SEA algorithm presented here adopts an approach similar to Clover, Pscan and AME. The input to SEA is one or two sets of sequences and a set of motifs, encoded as PWMs. If only one set of (primary) sequences is provided, SEA will construct a control set by shuffling each of the primary sequences. Unlike previous MEA algorithms, SEA computes highly accurate estimates of the statistical significance of the enrichment of each motif. It accomplishes this by setting aside a portion of each input sequence set for use in optimizing its scoring function, and then performs a single statistical test of enrichment for each motif. As we shall show, the estimates of statistical significance provided by the AME and CentriMo algorithms are conservative, which can cause potentially interesting enriched motifs to fail to pass the significance threshold.

## 2 Results

### 2.1 The SEA algorithm

SEA assumes that each primary sequence may contain zero or one occurrences (sites) of the motif (the so-called “ZOOPS” model [1]). Motif enrichment analysis will not be negatively affected if a primary sequence contains more than one occurrence of a motif. Unlike CentriMo, SEA does not rely on motif sites clustering near the centers of sequences, making it applicable to types of data (e.g., gene promoter sequences) where this is not the case.

SEA creates a Markov model of a user-specified order from the control sequences. SEA uses the Markov model in conjunction with the PWM when counting matches to the motif to further bias the search away from motifs that are mere artifacts of the lower-order statistics of the input sequences.

SEA reports accurate significance estimates for each motif’s enrichment in contrast to AME and CentriMo, whose motif enrichment significant estimates tend to be too conservative. SEA accomplishes this by choosing an optimal score threshold for each motif using a fraction of the primary and control sequences that it sets aside (holds out) and does not use during the significance estimation process.

Finally, SEA will work with user-specified (“custom”) alphabets, as will AME and CentriMo, but not the other algorithm studied here. This allows SEA to be applied to a wide range of motif enrichment analyses, including in epigenetically modified DNA or post-translationally modified proteins.

In the remainder of this paper, we describe the SEA algorithm in more detail and present experimental results comparing its performance with three widely-used motif enrichment algorithms. For the experimental comparisons, we consider motif enrichment in ChIP-seq datasets (transcription factor (TF) binding motifs). We validate the enrichment predictions using motifs derived using a completely independent assay—high-throughput SELEX [5].

SEA takes as input a set of primary sequences and an (optional) set of control sequences. In addition, SEA requires the user to provide one or more files containing known sequence motifs in the form of position weight matrices (PWMs). SEA computes the statistical significance of the enrichment of each motif in the motif dataset(s). SEA proceeds by first preparing the input sequence datasets and then by then computing the significance of each of the input motifs.

SEA first reads the input sequence dataset(s) (primary, and optionally, control), converting to upper-case if the sequence alphabet is not case sensitive, and converting all ambiguous characters to a “separator” character that is not present in the alphabet.

To ensure that SEA will give the same results regardless of the order of the sequences in the input dataset(s), it sorts the input dataset(s) alphabetically by sequence content, and then randomizes the order of the sequences in the dataset(s).

Next, if the user does not provide a set of control sequences, SEA creates one from the primary sequences. Each primary sequence is shuffled, preserving the frequencies of all words of length *k* (“*k*-mers”) within it, where *k* can be specified by the user. The shuffling also preserves the positions of any separator characters. This prevents artifacts that can be caused by the presence of ambiguous characters in the sequences (such as the “N” character used by DNA repeat-masking programs).

SEA chooses the statistical test it will use. It will use Fisher’s exact test if the primary and control sequences have the same average length (within 0.01%), otherwise it will use the Binomial test.

Then, SEA creates a Markov model of the control sequences of order *k* − 1. SEA uses this model in conjunction with the PWM to compute the likelihood ratio scores of words.

Next, SEA creates a “hold-out” dataset for accurately assigning statistical significance to the enrichment of each query motif. By default, the hold-out set consists of a random sample of 10% of the sequences in the primary and control datasets.

Finally, SEA computes the statistical significance of the enrichment of each motif. For each motif, SEA first uses it to score each of the primary and control sequences. Then, using just the hold-out sequences, it determines the score threshold that gives the optimal classification according to the chosen statistical test. Then using that score threshold, it classifies the remaining sequences and computes the statistical significance of the classification. This second significance value (*p*-value) is unbiased because it is based on only a single test per motif. Classification is based on the best match to the motif in each sequence (on either strand when the alphabet is complementable).

The SEA algorithm outputs its results in the form of an HTML file. A screenshot of a portion of an example HTML output file from SEA is shown in Fig. 1. The results are shorted in order of decreasing statistical significance and feature a sequence logo of the enriched motif, the name of the file (“Database”) containing the motif, its ID, various measures of statistical significance of the motif’s enrichment, counts of the number of times the motif occurs in the primary sequences (“TP”, true positives), or the control sequences, (“FP”, false positives), the positional distribution of the motif sites in the primary sequences, and a histogram of the number of motif sites per primary sequence. SEA also outputs its results in a tab-separated values file (TSV file) suitable for use with spreadsheet programs (e.g., Excel). If requested by the user, SEA will also output a list of the sequences that contain predicted sites for each enriched motif.

**Figure 1:**
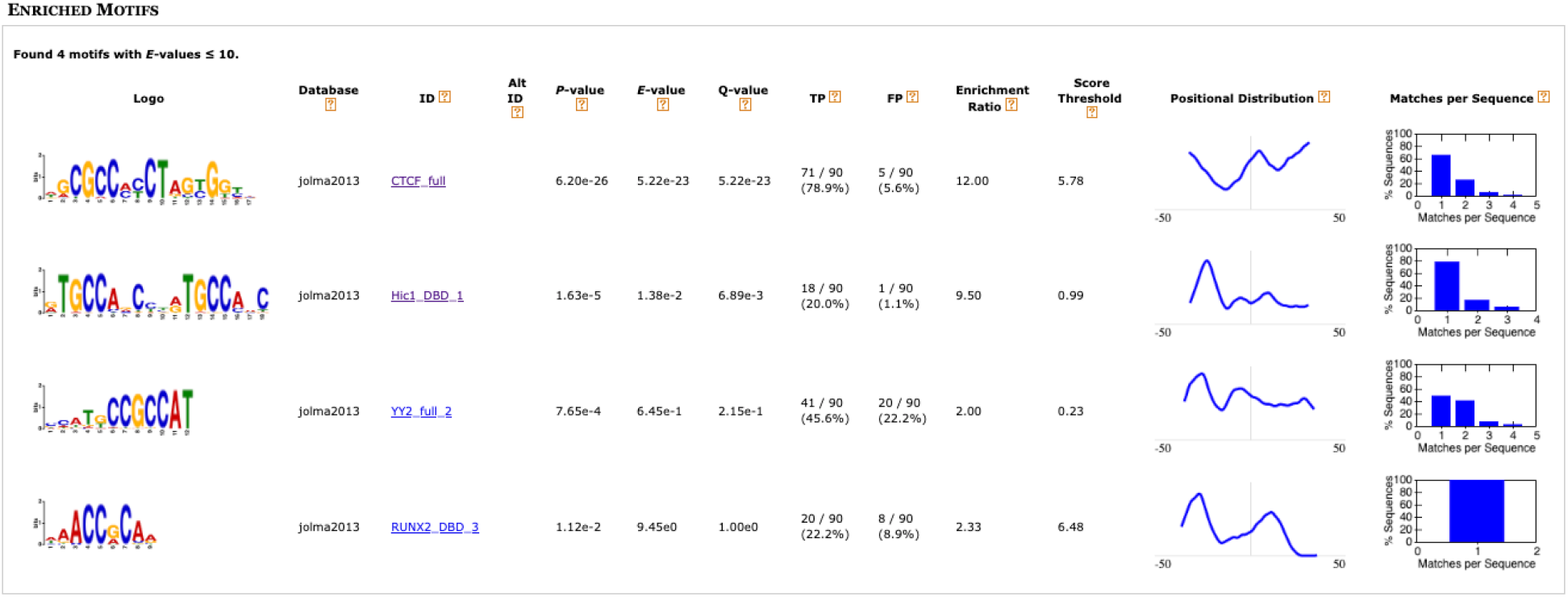
Screenshot of (a portion of) the HTML output of SEA.

### 2.2 Comparison of SEA with other MEA algorithms using TF ChIP-seq data

We compare SEA to three existing MEA algorithms: AME, CentriMo and Pscan. Our tests measure the ability of each MEA algorithm to correctly identify that a known TF-binding motif is enriched in a set of sequences bound to the TF. For the sequence sets, we use selected TF ChiP-seq datasets from the ENCODE consortium [2]. We selected all 44 datasets from experiments in K562 and Gm12878 cells where the ChIP-ed TF has a SELEX motif in the Jolma compendium [5]. In each of the following analyses, we run the MEA algorithms using as queries all 843 motifs in the Jolma compendium and determine where the algorithm ranks the known motif for the TF. We did not include the Clover algorithm in our tests because, with large sequence datasets, it runs extremely slowly compared with the SEA, AME, CentriMo and Pscan.

#### 2.2.1 Accuracy

One use case for MEA algorithms is predicting the identities of transcription whose binding sites are enriched in a given set of genomic sequences. To evaluate MEA algorithms on this task, we use TF ChIP-seq data, which we know should be highly enriched for the binding sites of the ChIP-ed TF. In this test, each MEA algorithm computes an enrichment score for each of the 843 SELEX motifs in the Jolma compendium, one of which is the motif for the ChIPed TF. Ideally, the enrichment of the known motif should be ranked highest (rank 1). We display the results as boxplots of the rank of the known motif in each of the 44 runs of an MEA algorithm in Fig. 2. (There is one run of each algorithm for each of the 44 ENCODE datasets).

**Figure 2:**
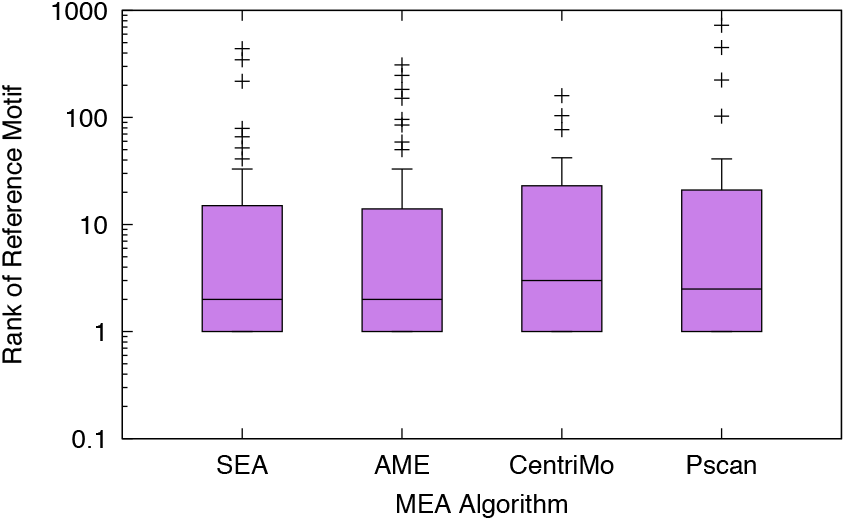
Accuracy of motif enrichment algorithms on ENCODE TF ChIP-seq datasets. Boxplots show the distribution of the rank of the reference motif for the ChIP-ed TF in the output of the named MEA algorithm. Each algorithm is run once using each of 44 ENCODE ChIP-seq datasets, with 843 SELEX query motifs (including the known motif). The boxplots show the range of the middle quartiles with a line at the median, and stars show outliers farther than 1.5 times the interquartile range (the whiskers) from the median.

It is evident in Fig. 2 that the four MEA algorithms compared here have similar accuracies at ranking motifs for enrichment in this test. The median rank of the reference motif for the ChIP-ed TF is 2 for both SEA and AME, and only slightly worse for CentriMo and Pscan. The overall distributions of the predictions are also similar.

#### 2.2.2 Sensitivity

To further explore the performance of the MEA algorithms on the task of correctly identifying enriched motifs, we modified the ChIP-seq datasets to make the MEA task more difficult in two ways. Firstly, we compare the ability of MEA algorithms to correctly determine the identity of the ChIP-ed TF as a function of the amount of noise in the sequences. To this end, we add varying amount of noise to each of the 44 ENCODE ChIP-seq datasets by shuffling the letters in a fraction of the sequences. We say that the original dataset has 100% purity, and that a dataset where 99.5% of the sequences have been shuffled has 0.5% purity. Secondly, we compare the ability of the algorithms to determine the ChIP-ed TF as a function of the total number of sequences in their input. We do this by sampling varying numbers of sequences from each of the 44 ChIP-seq datasets.

We say that an algorithm succeeds for a ChIP-seq dataset if it assigns the SELEX motif for the ChIP-ed TF the *highest* significance among all the 843 SELEX motifs in the Jolma2013 compendium. This is a very strict criterion. An algorithm that ranks the reference motif second is considered to have failed by this metric. Fig. 3 shows the results of these comparisons.

**Figure 3:**
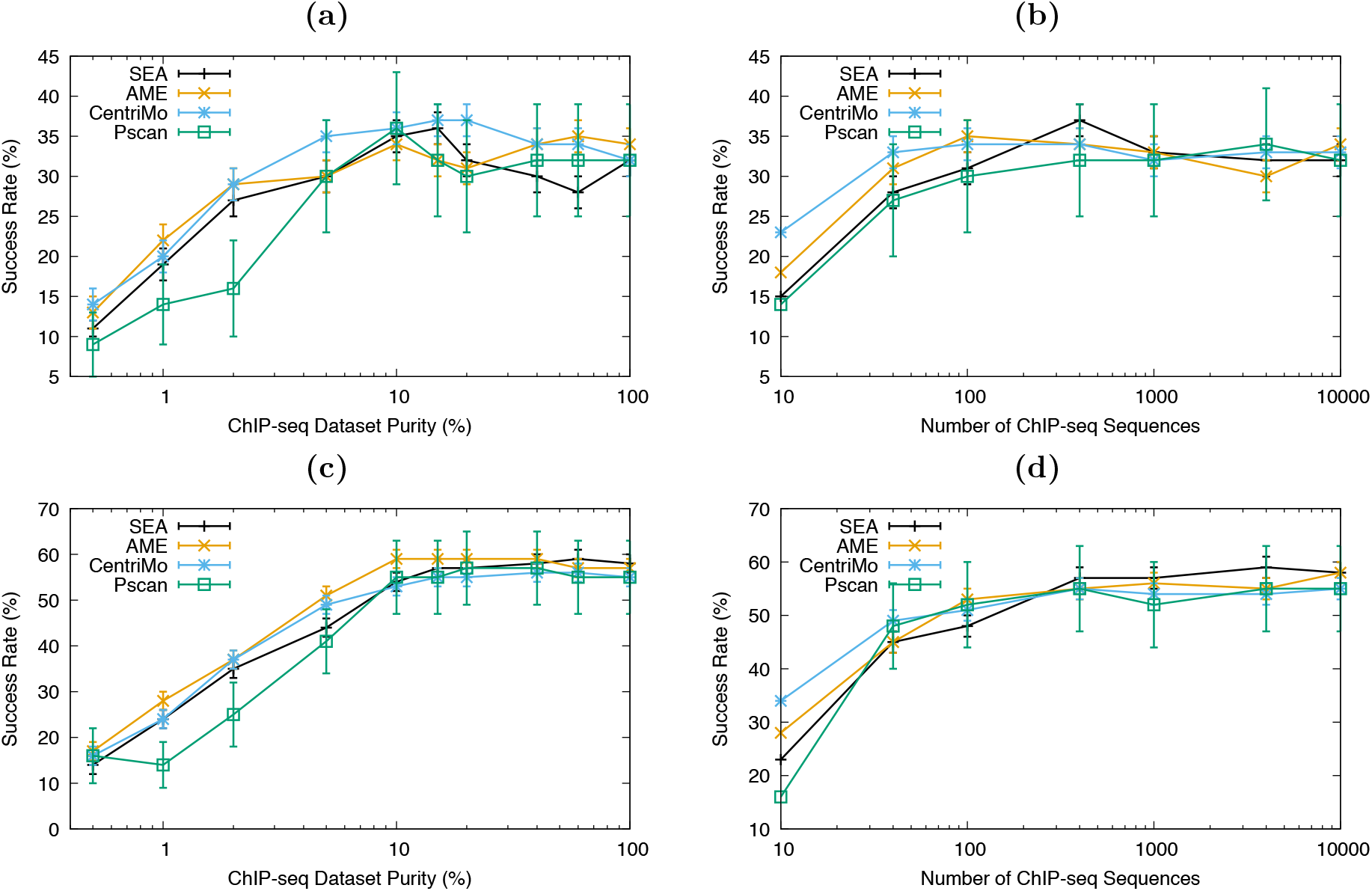
Sensitivity of MEA algorithms on ENCODE TF ChIP-seq datasets. The curves show the percentage of times (*Y*) the known motif has rank 1 in the output of the given MEA algorithm, averaged over 44 ChIP-seq datasets as a function of the purity (Panel **a**) or the size (Panel **b**) of the datasets. Panels **c**) and **d**) present equivalent results for the percentage of time the known motif is ranked among the top 3 by the MEA algorithm.

We observe in Fig. 3a that the four MEA algorithms compared here perform quite similarly to each other as we add increasing amounts of noise to the input sequences. At most noise levels, the success rate of the algorithms fall within each others error bars. Pscan may perform slightly worse than SEA, AME and CentriMo at higher level of noise.

Similarly, Fig. 3b shows that the four algorithms mostly have comparable performance when we reduce the number of ChIP-seq sequences. We do notice a difference when there are fewer than 100 sequences in the input set, with CentriMo having a higher success rate at ranking the reference motif highest among the 843 input motifs.

The above observations remain true even when we relax the criterion for success to ranking the reference motif among the top 3 input motifs.

#### 2.2.3 Speed

Fig. 4 shows the running time of the MEA algorithms on each of the 44 experiments described above in the Accuracy section. We run the algorithms using a single thread on a 3.2 GHz Intel Core i7 processor with 16GB of memory. We display the results as boxplots of the running times of each of the MEA algorithms on the 44 ChIP-seq datasets. It is apparent from Fig. 4 that the SEA algorithm is the fastest of the four that we test here, and Pscan is by far the slowest.

**Figure 4:**
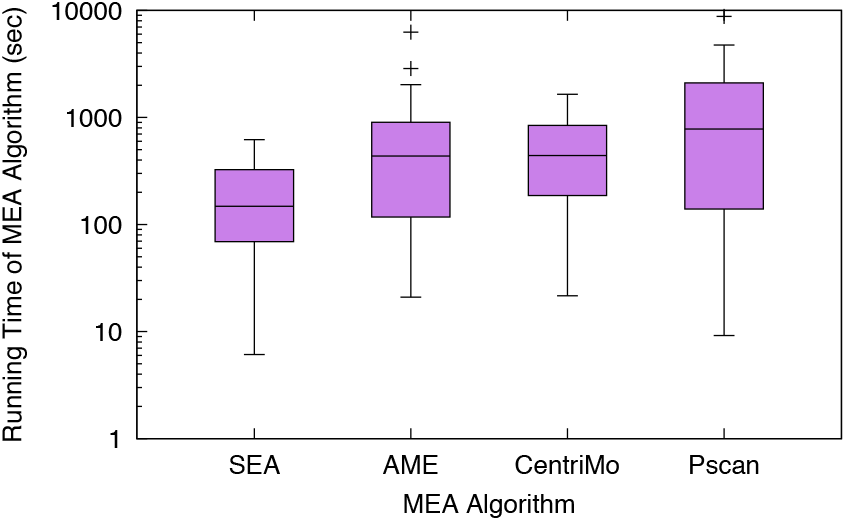
Speed of motif enrichment algorithms on ENCODE TF ChIP-seq datasets. Boxplots show the distribution of the running time of the named MEA algorithm. Each algorithm is run once using each of 44 ENCODE ChIP-seq datasets, with 843 SELEX query motifs. The boxplots show the range of the middle quartiles with a line at the median, and stars show outliers farther than 1.5 times the interquartile range (the whiskers) from the median.

### 2.3 SEA accurately estimates motif enrichment significance

The MEA algorithms studied here all report estimates of the statistical significance of the enrichment of each query motif. To verify the accuracy of these *p*-values, we run each MEA algorithm on 100 randomly generated datasets, each containing 1000 sequences. Since the sequences are random, the *p*-values for all the query motifs should follow a uniform distribution. To check that they are uniformly distributed, we create a Q-Q plot [10], which plots the theoretical value of the *n*th largest *p*-value, *x* = 1/(*n* + 1), versus the *p*-value reported by the MEA algorithm, *y*.

The *p*-values estimated by SEA using random sequences are highly accurate, as seen in Fig. 5, consistently lying within a factor of 2 of the theoretical uniform distribution. This is true with DNA, RNA and protein sequences. By contrast, AME and CentriMo *p*-values are consistently conservative (too large), often by over an order of magnitude. In practice, this can cause truly enriched motifs to be rejected. The *p*-values reported by Pscan do not seem to be adjusted correctly for multiple testing, as they are consistently too small. This makes it impossible to use Pscan *p*-values to determine if a motif is significant or not.

**Figure 5:**
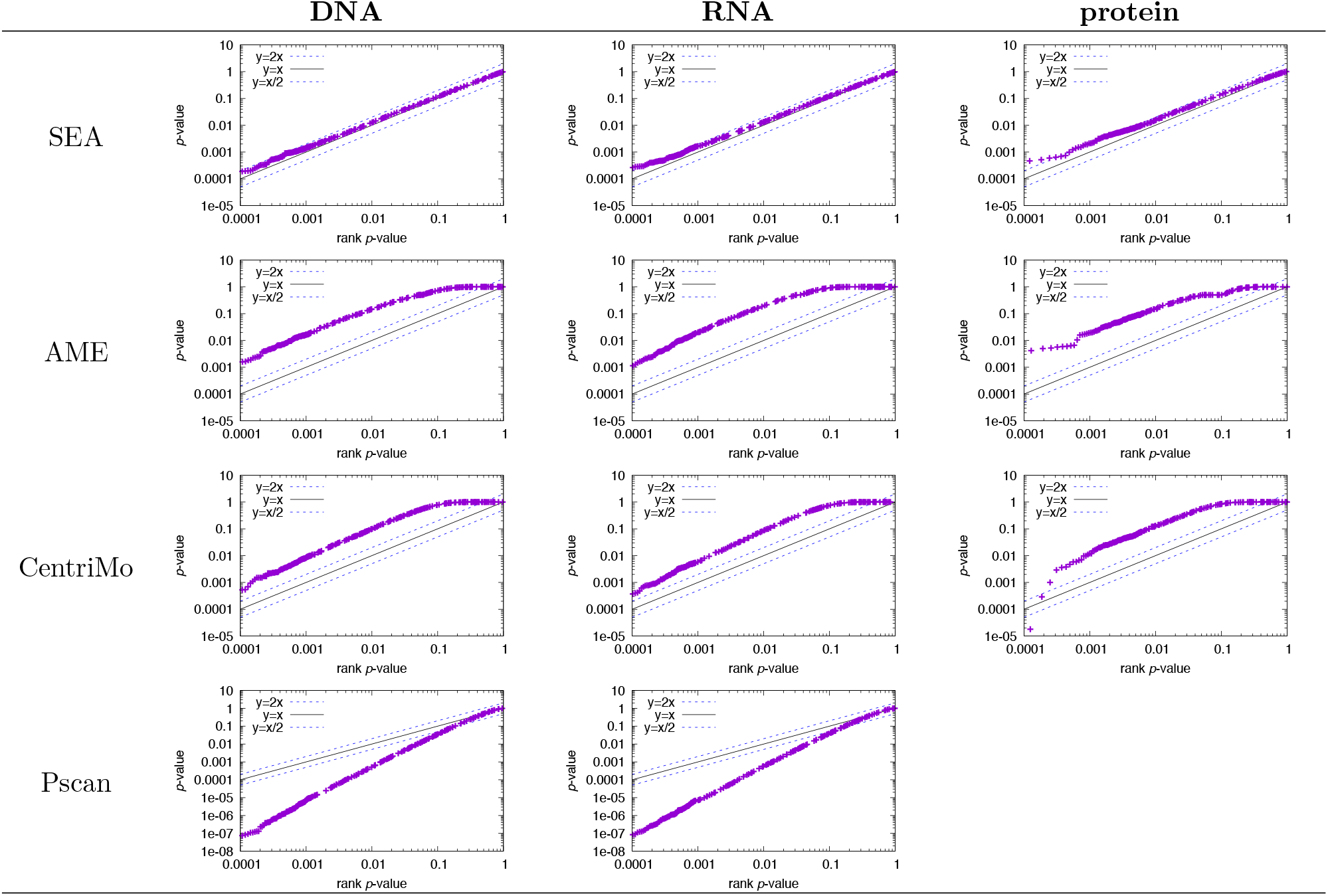
Q-Q accuracy plots of the *p*-values reported by different MEA algorithms. Each panel shows the Q-Q plot for the *p*-values reported by the named MEA algorithm when run on 100 datasets containing 1000 random sequences over the alphabet given above the panel. The sequences are 100 characters long for SEA, AME, and Pscan, and 500 characters long for CentriMo. Pscan does not accept protein sequences. Ideally, the points should lie along the line *Y* = *X*.

## 3 Methods

### 3.1 SEA algorithm details

#### 3.1.1 Statistical tests of motif enrichment

When the primary and control sequences have identical length distributions, SEA uses the Fisher exact test to determine if a motif is enriched. Suppose that there *N_p_* and *N_c_* primary sequences and control sequences, respectively, and there are sites in *n_p_* and *n_c_* of the primary and control sequences, respectively. The Fisher exact test assumes a null model where a site is equally likely likely to be in any sequence. It computes the probability that *n_p_ or more* primary sequences would contain a site, and *n_c_ or fewer* control sequences would contain a site, given the values *N_p_*, *n_p_*, *N_c_* and *n_c_*.

When the primary and control sequence sets have different length distributions SEA uses the Binomial test (rather than Fisher’s exact test) to estimate the statistical significance of the enrichment of a query motif. Suppose that there are *N_p_* primary sequences with average length *L_p_*, and *N_c_* control sequences with average length *L_c_*. Then, on average, the number of possible positions a motif of width *w* could occupy is approximately *S_p_* = *N_p_*(*L_p_* − *w* − 1) and *S_c_* = *N_c_*(*L_c_* − *w* − 1), respectively. So SEA estimates the Bernoulli probability *P_b_* that a site chosen randomly in either of the two sets of sequences actually is in a primary sequence as *P_b_* = *S_p_/S_c_*. If *n_p_* primary sequences contain a match to a given motif (*n_p_* “successes”), the Binomial test considers the statistical significance of the motif to be the probability that *n_p_* or more primary sequences have matches to the motif. The test estimates the *p*-value of the motif as the sum of the binomial probabilities of *k* successes in *N_p_* trials, each with probability of success *P_b_*, where *k* goes from *n_p_* to *N_p_*,

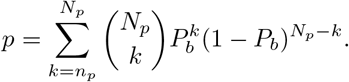

#### 3.1.2 Scoring sequences with motifs

SEA computes the log-likelihood score of each position in each primary and control sequence. If the Markov background model specified by the user is 0-order, SEA converts the motif frequencies into a standard PWM. This PWM records the log-likelihood ratio for each possible letter at each position in the motif,

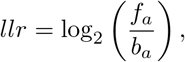

where *f_a_* is the frequency of letter *a* according to the motif, and *b_a_* is the frequency of the letter according to the background model. If the user specifies a higher-order background model, SEA sets *b_a_* = 1. To score a word in a sequence, SEA adds the entries in the PWM corresponding to the letters in the word. If the background model is higher-order, SEA the sub-tracts the logarithm of the probability of the word given the background model. For efficiency, SEA pre-computes the cumulative probability of each position in the sequence according to the (higher-order) background model.

### 3.2 Evaluating motif enrichment analysis algorithms with TF ChIP-seq datasets

#### 3.2.1 Reference motifs

In order to evaluate the accuracy, sensitivity and speed of motif enrichment analysis algorithms on ChIP-seq datasets, we identify 22 ENCODE TF ChIP-seq experiments in K562 cells and 22 experiments in Gm12878 cells for which there is a motif derived from high-throughput SELEX data for the same TF in the Jolma compendium of 843 motifs [5]. We download the 44 ENCODE AWG narrowPeak BED files of TF ChIP-seq experiments from the UCSC server at http://hgdownload.cse.ucsc.edu/goldenPath/hg19/encodeDCC/wgEncodeAwgTfbsUniform. For some TFs there are multiple SELEX motifs in the compendium corresponding to full-length proteins (names ending in `_full`) or DNA-binding domains (`_DBD`). In addition, the compendium sometimes contains multiple, alternative motifs for full-length proteins and/or DNA-binding domains. We arbitrarily choose the lowest-numbered of these. Thus, we choose as the reference motif for a ChIP-seq dataset the SELEX motif whose name matches the ChIP-ed TF name and ends in 1) `_full`, 2) `_full_1`, `_DBD` or 4) `_DBD_1`, in order of preference. The names of the 44 ENCODE AWG BED files and the names of their reference motifs (which are available at http://meme-suite.org/meme-software/Databases/motifs/motif_databases.12.19.tgz in file `EUKARYOTE/Jolma2013.meme`) are given in Table 1.

**Table 1:**
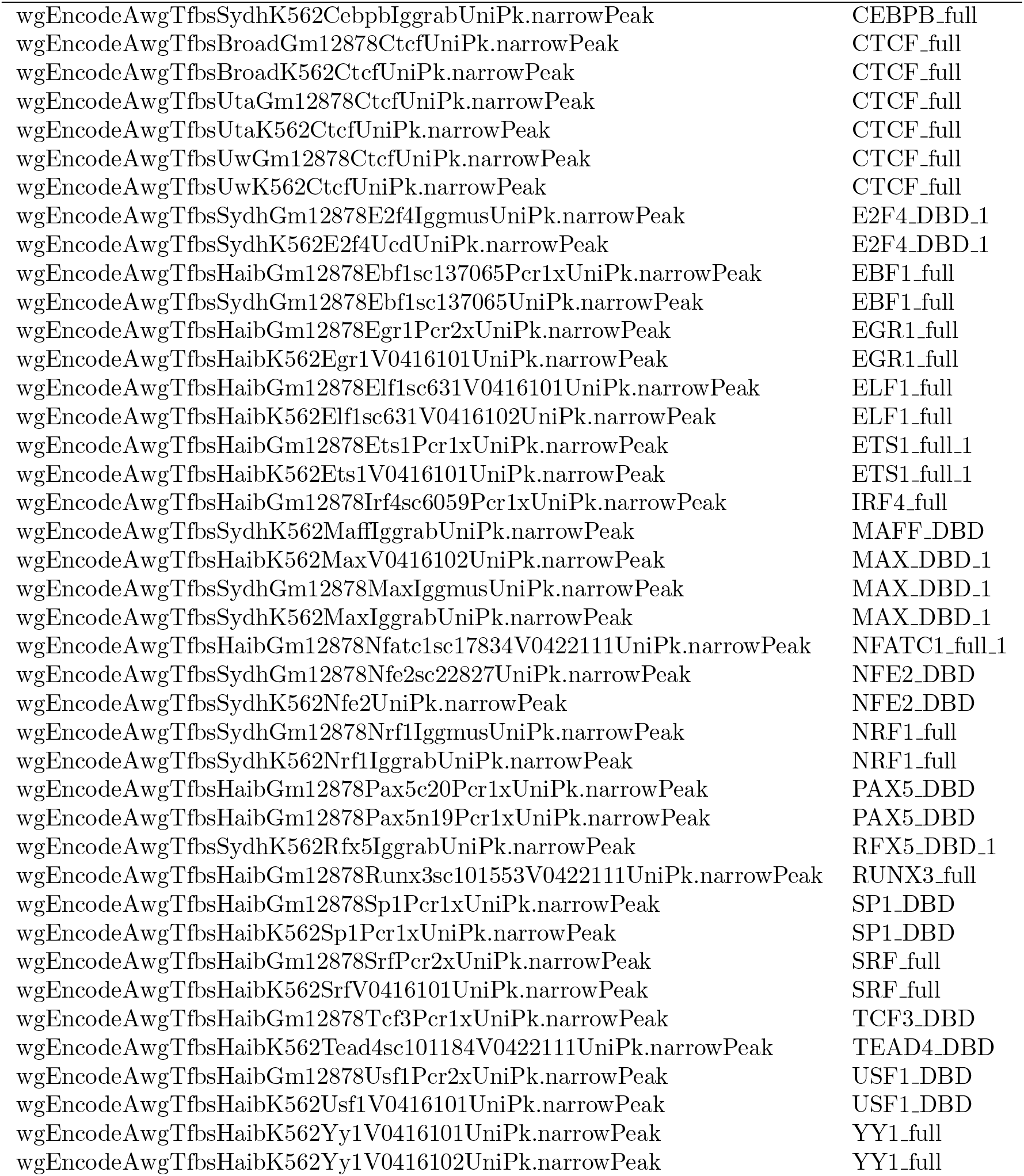
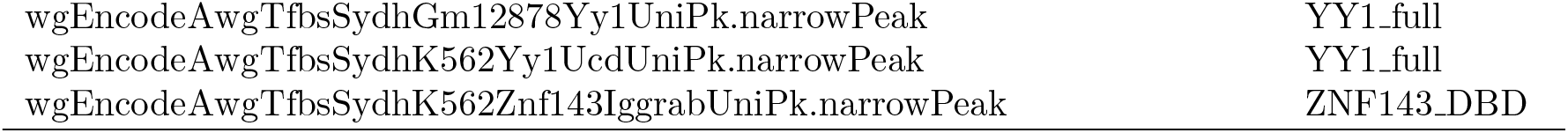
ENCODE K562 and Gm12878 TF ChIP-seq files and SELEX [5] reference motifs. ChIP-seq files are available at http://hgdownload.cse.ucsc.edu/goldenPath/hg19/encodeDCC/wgEncodeAwgTfbsUniform with extension gz. Motifs are available at http://meme-suite.org/meme-software/Databases/motifs/motif_databases.12.19.tgz in file `EUKARYOTE/Jolma2013.meme`.

#### 3.2.2 Primary and control sequence datasets

For each of the 44 ENCODE narrowPeak files, we create a FASTA files of 100bp sequences and 500bp sequences, respectively, centered on each of the peaks. We use the `fasta-fetch-centered` tool available in the MEME Suite software package to extract sequences from the *Homo sapiens* genome (hg19), which we download from the UCSC genome browser website (http://hgdownload.soe.ucsc.edu/goldenPath/hg19/bigZips/hg19.fa.gz). We use these files, which contain between 772 and 67,417 sequences, as the primary sequence datasets. The 500bp sequences are used with CentriMo, and the 100bp sequences are used with the other MEA algorithms. For each 100bp dataset, we create a control dataset by shuffling each sequence in the primary dataset using the `fasta-shuffle-letters` tool available in the MEME Suite software package. With this tool, the user can specify that the frequencies of words (*k*-mers) of any size *k* be preserved. The control dataset sequences are thus 100bp long, and have lower-order statistics matching those of the corresponding primary dataset.

#### 3.2.3 Evaluating algorithm performance

To evaluate the accuracy and sensitivity of the MEA algorithms, we use each algorithm to estimate the enrichment of the 843 motifs in the Jolma compendium in the input sequences corresponding to a particular ChIP-seq experiment, and note the rank algorithm assigns to the reference motif for that experiment (where rank 1 is the most significant motif). For the accuracy experiments, we do this once for each ChIP-seq experiment. For the sensitivity experiments, we do it ten times using ten different random seeds for choosing which sequences to shuffle (“purity”), or which which sequences to include in the primary dataset (“size”).

### 3.3 Evaluating p-values

To evaluate the accuracy of the *p*-values estimated by MEA algorithms, we use randomly generated sequences and motifs. We generate random DNA and protein sequence datasets using the gendb tool from the MEME Suite package. This tool allows us to specify the number as well as the minimum and maximum length of generated sequences. We generate RNA sequences by replacing T with U in DNA sequences generated by gendb.

To generate random motifs, we shuffled the columns of each motif in databases of DNA, RNA and protein motifs, respectively using the `meme-shuffle-motifs` tool from the MEME Suite package. We use the randomly generated sequences and the shuffled motifs as input to the the MEA algorithm, and record the *p*-value of the most significant (random) motif. We repeat this 1000 times, using a different random seed for shuffling the motif columns each time, and plot the resulting values as a Q-Q plot. The DNA, RNA and protein motif databases we shuffle are the Jolma compendium [5], CISBP2 [9], and ELM2018 [6], respectively.

## 4 Discussion

Our results show that SEA is a versatile new algorithm for analysis of the enrichment of sequence motifs. The main advantage of the SEA algorithm is the fact that it produces highly accurate estimates of the statistical significance (*p*-value) of motif enrichment, whereas the other three algorithms studied here produce less reliable *p*-values. The AME and CentriMo estimates are conservative by an order of magnitude or more, which can cause truly enriched motifs to fail to pass a chosen *p*-value threshold. In contrast, Pscan produces grossly liberal estimates that can not safely be used to judge motif enrichment. Another advantage of the SEA algorithm is that it is faster than AME and CentriMo, and much faster than Pscan.

The other MEA algorithms studied here have their own advantages. AME is primarily designed to be used when the user has additional information in the form of a number (e.g, fluorescence intensity from a microarray experiment) about each of the input sequences. AME utilizes this information to rank the sequences and measures motif enrichment in terms of the correlation between the auxiliary number and the motif score of each sequence. CentriMo is only designed for use when the the motifs tend to be located near the center of the input sequences (e.g., ChIP-seq peaks). We note that CentriMo was more sensitive than the other algorithms when the number of ChIP-seq peaks in its input was very small (fewer than 100). Finally, Pscan is very convenient to use to identify motifs enriched in sets of promoters from the six model organisms supported by its website (http://www.beaconlab.it/pscan).

